# Essential conserved neuronal motors kinesin-1 and kinesin-3 regulate Aβ42 toxicity in vivo

**DOI:** 10.1101/2024.04.23.590704

**Authors:** Deepthy Francis, Francesco Paonessa, Caroline C.G. Fabre, Victoria L. Hewitt, Maria E. Giannakou, Isabel Peset, Alexander J. Whitworth, Frederick J. Livesey, Isabel M. Palacios

## Abstract

Alzheimer’s Disease is the leading cause of dementia and the most common neurodegenerative disorder. Understanding the molecular pathology of Alzheimer’s Disease may help identify new ways to reduce neuronal damage. In the past decades *Drosophila* has become a powerful tool in modelling mechanisms underlying human diseases. Here we investigate how the expression of the human 42-residue β-amyloid (Aβ) carrying the E22G pathogenic “Arctic” mutation (Aβ_42Arc_) affects axonal health and behaviour of *Drosophila*. We find that Aβ_42Arc_ flies present aberrant neurons, with altered axonal transport of mitochondrial and an increased number of terminal boutons at neuromuscular junctions. We demonstrate that the major axonal motor proteins kinesin-1 and kinesin-3 are essential for the correct development of neurons in *Drosophila* larvae and similar findings are replicated in human iPSC-derived cortical neurons. We then show that the over-expression of kinesin-1 or kinesin-3 restores the correct number of terminal boutons in Aβ_42Arc_ expressing neurons and that this is associated with a rescue of the overall neuronal function, measured by negative geotaxis locomotor behavioural assay. We therefore provide new evidence in understanding the mechanisms of axonal transport defects in Alzheimer’s Disease, and our results indicate that kinesins should be considered as potential drug targets to help reduce dementia-associated disorders.

## Introduction

Alzheimer’s Disease (AD) is the leading cause of dementia and the most common neurodegenerative disorder. Age is a major risk factor for AD, but genetic studies show that the amyloid precursor protein (APP) and its processing play a critical role in AD^1,2^. Neurons process APP through the amyloidogenic pathway where the protein is sequentially cleaved by β-secretase (<u>b</u>eta site <u>A</u>PP <u>c</u>leaving <u>e</u>nzyme BACE) and γ-secretase (with the catalytic subunit presenilin). Processing by these two enzymes generates the peptide Aβ_1-40_ and the more toxic peptide Aβ_1-42_ (Aβ_42_ from hereon). In Alzheimer’s disease and other dementias, Aβ_42_ oligomerises and forms aggregates in the extracellular space, which ultimately form the macroscopic plaques that are a main hallmark of AD pathology. This deposition of extracellular Aβ oligomers has been widely associated with detrimental effects on cellular physiology, such as disruption of the microtubule cytoskeleton^3,4^ and impairment of axonal transport^5–8^. In the last two decades, the fruit fly *Drosophila melanogaster* has been used as a model for AD studies, where two different Aβ proteotoxicity models have been investigated: APP/presenilin/BACE-expressing transgenic flies, which exhibit age-dependent AD-like pathology and behavioural changes as a consequence^9–14^, or human Aβ expressing flies. One specific fly model relies on the human Aβ_42_ with the “Arctic” E22G mutation (Aβ_42Arc_ from hereon), which causes an aggressive form of familial AD^15^. The expression of Aβ_42Arc_ in this humanised *Drosophila* line results in a severe form of neurodegeneration, associated with progressive locomotor deficits, brain vacuolation and premature death of the animals^9,16^.

It has been argued that Aβ may also affect neuronal physiology through the disruption of intracellular processes, for example by disrupting microtubule-based axonal transport^17,18^. Neurons are critically dependent on intracellular transport of components and genetic mutations in the transport machinery are linked to axonopathy and neurodegeneration^19,20^. Axonal transport is facilitated by microtubule motors, with kinesin-1 (KIF5 motors in mammals, and kin1 from hereon) and kinesin-3 (KIF1 motors in mammals, and kin3 from hereon) mediating anterograde transport from the cell body to distal parts of the axon of various cargoes such as mitochondria and vesicles. Recent work suggests a close association between Aβ toxicity and the loss of the mammalian neuronal kin1 KIF5A^21^, that is likely the major motor for APP vesicle transport^18,22–24^. However, it is unknown if increasing the activity of transport kinesins, such as kin1 and kin3, leads to a rescue of AD symptoms. Since axonal transport and Aβ toxicity are conserved in humans and flies, a better understanding of these mechanisms using humanised *Drosophila* could help with human studies^25–27^. The reason for using humanised flies is that it constitutes a cost-effective, fast model system with powerful genetics to address mechanistic questions. In addition, key proteins in our study, such as kinesins, are encoded by a single gene in *Drosophila*, contrary to mammals, facilitating functional analysis and gene manipulations.

In this work, we used the humanised Aβ_42Arc_ flies and human iPSCs-derived neurons to further explore the link between pathological Aβ peptides, axonal transport, and neurodegeneration. We demonstrate that the depletion of kin1 and kin3 from motor neurons (MNs) in *Drosophila* larvae led to their death, and we obtained similar results in human iPSC-derived cortical neurons depleted of the kin1 and kin3 ortholog genes. Expression of the pathologic form of human Aβ_42Arc_ in *Drosophila* MNs inhibits mitochondrial transport in the distal axons. The flies expressing Aβ_42Arc_ in MNs also showed aberrant neuromuscular junction (NMJ) synapses, with increased number of terminal boutons, a phenotype which was rescued by the over-expression of kin1 and kin3. Flies expressing Aβ_42Arc_ in MNs also showed reduced climbing activity, a typical behaviour linked to neuronal health, and used as a common read-out for neuronal toxicity in flies. We observed a rescue of this phenotype when kin1 or kin3 were over-expressed in the same neurons. We propose that modulation of kin1 and kin3 activity may represent a promising target for pharmacological intervention in AD.

## Results

### kin1 and kin3 are essential for survival of motor neurons in *Drosophila melanogaster*

To study the relationship between kinesins and neurodegeneration, we first analysed whether kin1 and kin3 are required for neuronal survival. We obtained larvae with a subset of MNs lacking kin3 (*imac^1^*^70^ allele with the mutation W58term^28^) or lacking the motor subunit of kin1 (*Khc^27^*null allele^29^). This was achieved by clonal induction using the MARCM system (Mosaic Analysis with a Repressible Cell Marker^30^), so that any mutant MN lacking kin1 or kin3 is labelled with myristoylated-RFP (myr-RFP, a lipid-modified reporter protein). Our MARCM approach allows for the cell body, dendrites, axons and NMJs of the mutant MNs to be visualised in live animals (Supplementary Figure S1). The viability of those mutant MNs was then monitored as the larvae developed from second instar larvae L2 (∼first instar larvae L1+24h) to third instar larvae L3 (∼L2+48h) by quantifying the number of L2 animals exhibiting mutant myr-RFP positive neurons, and then screening the same animals for the presence of those neurons at L3 stage (Figure 1). Quantification of larvae containing *Khc*^27^ MNs shows that only 37% (n=23; p<0.0001) of the larvae displayed myr-RFP positive *Khc*^27^ mutant MNs at the L3 stage, compared to the L2 stage (Figure 1B). For larvae with *imac*^17^ MNs, 73% (n=22; p<0.001) of the larvae displayed myr-RFP positive *imac^1^*^70^ mutant neurons at the L3 stage, compared to the L2 stage (Figure 1B). Thus, a significant number of mutant neurons present at stage L2 were lost during the L2-to-L3 development of the larvae. No loss of induced mutant clones over time was observed in the control clones (n=27, controls are MARCM clones without any *kin* mutation associated). These experiments show that in the developing nervous system, neurons lacking kin1 or kin3 disappear as the larvae grow from L2 to L3. This finding indicates that kin1 and kin3 are essential for the survival of developing MNs.

**Figure 1.**
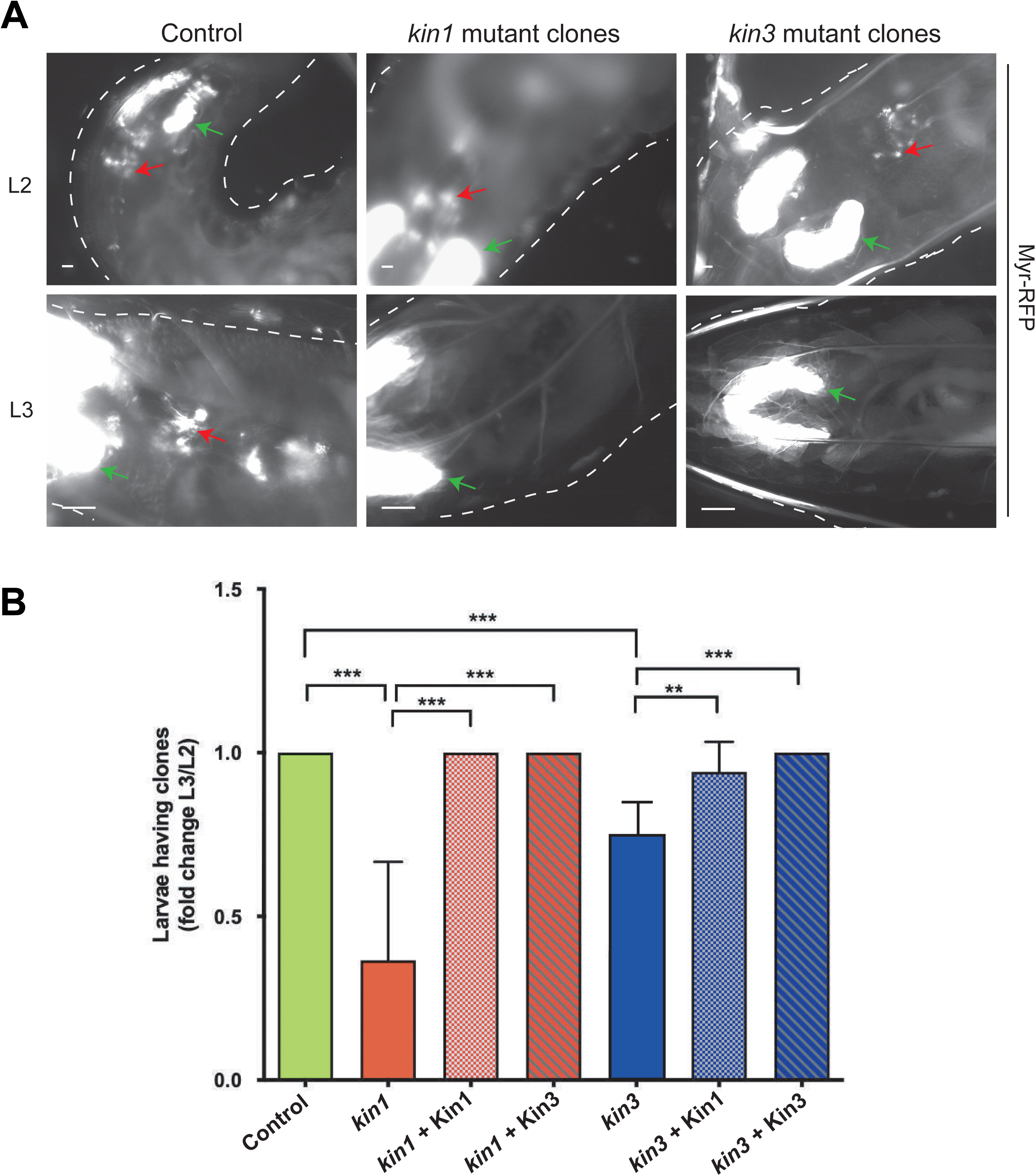
kinesin-1 and kinesin-3 are required for survival of neuronal clones in *Drosophila* larvae. (A) Analysis of MARCM-induced clones of *Drosophila* larvae at the 2^nd^ (L2 (∼L1+24h)) and 3^rd^ instar (L3) (∼L2+48h) in the same individual, where clones (red arrows) in the motor neurons (driven by the OK371-Gal4 driver line) are marked with myristoylated-RFP, which labels all the membranes of the cell body, axons and dendrites. The larvae are oriented with the anterior region towards the left and the posterior region towards the right. The anterior region of each larva can be distinguished by the presence of two bright salivary glands (green arrows); the CNS is located in between the two salivary glands. Representative images are shown from FRT42D control (left), *kin1* mutant clones (middle), and *kin3* mutant clones (right) in the region of the salivary glands. Note that in the control, both L2 and L3 (of the same individual) show several clones that are visible as marked with RFP (red arrows). However, analysis at L3 of *kin1* mutant and of *kin3* mutant larvae shows loss of clones that were present at L2 (red arrows). Scale bar, 100μm and 300μm for L2 and L3, respectively. (B) Histogram showing the average percentage of animals displaying MARCM-induced clones from L2 to L3 (same animals) in control (n=27), *kin1* (n=23) and *kin3* (n=22) mutant larvae, as well as in larvae carrying the *kin1* mutation in conjunction with Kin1 (n=20) or Kin3 (n=13) overexpression, and in larvae carrying the *kin3* mutation in conjunction with Kin1 (n=24) or Kin3 overexpression (n=19). *kin1* and *kin-3* mutant larvae show a significant loss of neuronal clones from L2 to L3, in comparison with the other genotypes analysed. Error bars represents ± SEM. * indicates p<0.01.

We then investigated whether re-expressing kin1 or kin3 in *kin1* (i.e., *Khc*^27^) and *kin3* (i.e., *imac^1^*^70^) mutant neurons could restore their viability. This was achieved by driving the expression of UAS-kin1-GFP or UAS-kin3-GFP in the MARCM clones (Figure 1B). We observed that the expression of Kin1-GFP or Kin3-GFP in *kin1* or *kin3* mutant clones, respectively, showed a complete elimination of neuronal death (100%, n=24 for kin1 and 100%, n=19 for kin3). Surprisingly, over-expression of Kin1-GFP in *kin3* mutant neurons eliminated neuronal death in 91.6% of the animals (n=24), and over-expression of Kin3-GFP in *kin1* mutant neurons also eliminated death in 100% of the animals (n=19), suggesting that motors may act in a redundant or concerted manner, and supporting the recent findings that the concerted action of a kin1 and a kin3 promotes efficient secretory vesicle transport^31^.

### Depletion of kin1 and kin3 represses the development of human iPSC-derived cortical neurons

To decipher whether kin1 and kin3 are also essential for neuronal survival in humans, we investigated the role of these two kinesins in the development of human iPSC-derived cortical neurons. For this purpose, we generated human cortical neurons from a control iPSC line using a well-established protocol^32^. We initially characterised the expression and localisation of KIF5A (human ortholog of kin1) and KIF1A (human ortholog of kin3 and a major mammalian neuronal kinesin) in mature iPSC-derived cortical neurons. Immunohistochemistry showed that KIF1A and KIF5A are expressed in human neurons and are found in processes with punctate pattern suggestive of cargoes (Figure 2A).

**Figure 2.**
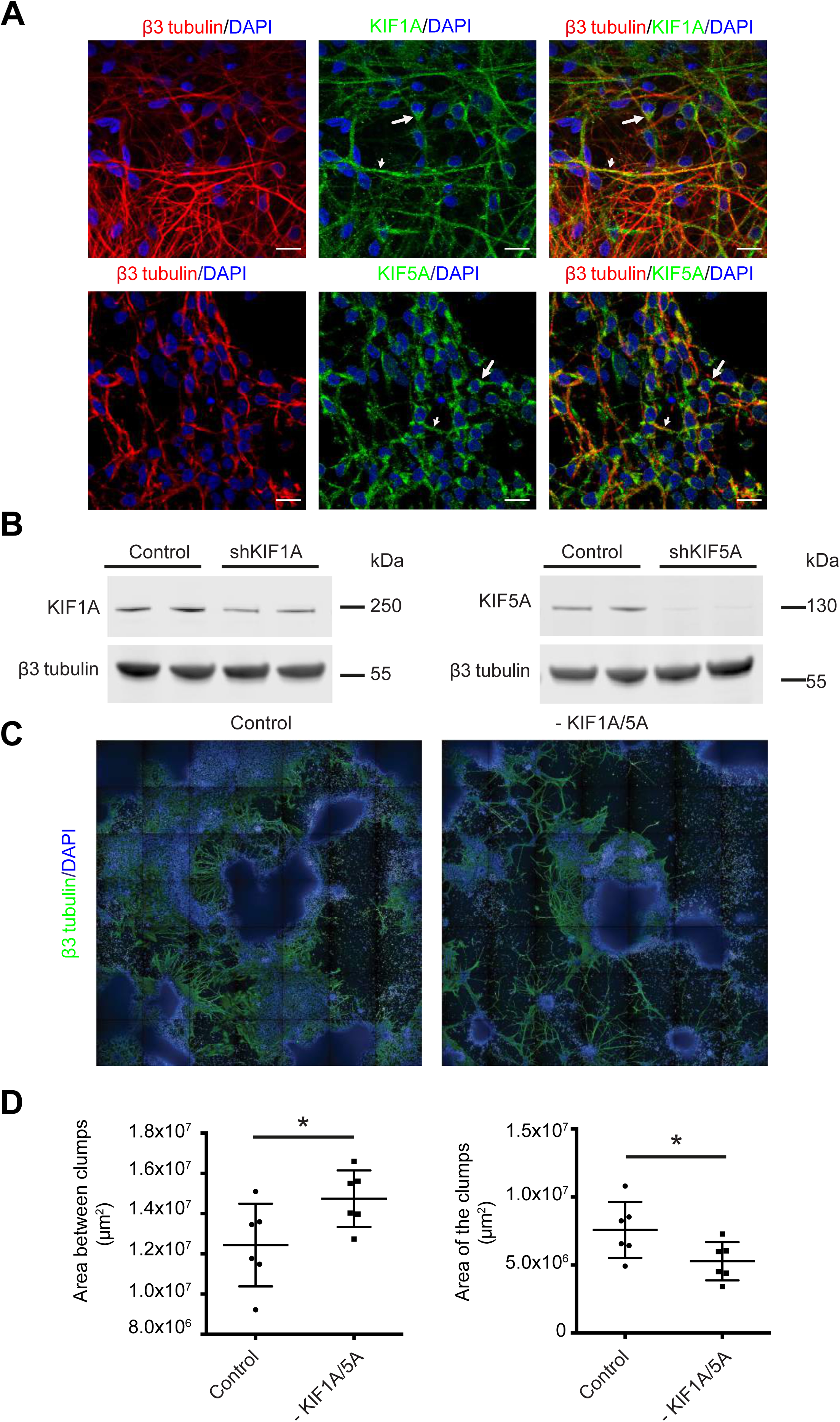
Silencing of KIF1A and KIF5A in iPSC-derived neurons. (A) iPSC-derived neurons were cultured for 100 days and then labelled using DAPI (blue), and antibodies against KIF1A (upper row; green) or KIF5A (bottom row; green) and β3-tubulin (labels axons and dendrites; red). Both motors are detected in cell body (long arrow) and neurites (short arrow), revealing a speckled pattern. Scale bars, 20μm. (B) iPSC-derived neurons were infected with scramble shRNA (control) or shRNA against KIF1A and KIF5A (see materials and methods). Western blots show the reduction in KIF1A and KIF5A proteins after shRNA infection. The untargeted β3-tubulin was not affected. (C) iPSC-derived neurons were infected at day 30 with both shRNA constructs against KIF1A and KIF5A and cultured to day 65 when cells were fixed and stained for DAPI (blue) and β3-tubulin (green). The composite images capture cell distribution in the entire imaging well of a 96 well plate. It is possible to notice a reduction of both cell growth and neurites extension in neurons with shRNAs against KIF1A and KIF5A, compared with the control (scramble shRNA). (D) Cell growth was analysed trough the Harmony software using DAPI area as a marker of cell distribution. We observe an increase of the area between DAPI clumps and a reduction of clump area in neurons infected with shRNA against KIF1A and KIF5A (n=6); Student’s t test (* indicates p<0.05). Lines show means and SEM.

Next, we wanted to test whether kinesins are required for the correct maturation of human cortical neurons. In healthy conditions, iPSCs give rise to neural progenitor in 25 days. Progenitor cells initially generate cortical rosettes, a structure morphologically like the developing neural tube, that gradually expand their size becoming visible clumps that generate post-mitotic cortical neurons able to develop an intricate and functional network^33^. We thus measured the ability of cortical progenitors to form clumps and generate a complex neuronal net in absence of kinesins. Progenitor cells were infected with lentiviral vectors encoding shRNAs for KIF1A and KIF5A, reducing the expression of either (sh KIF1; sh KIF5A) or both (sh KIF1A/5A) motor proteins (Figure 2B-D; Supplementary Figures S2 and S3). Sh KIF1A/5A cells showed a dramatic decrease of the size of neuronal clumps when compared to control, as well as a reduction in the area covered by neuronal processes, highlighted by the presence of large void areas between neurogenic clumps (Figure 2C-D). This was confirmed by quantification, showing fewer cells in areas between neuronal clumps (Supplementary Figure S2A-B). This suggests that a simultaneous depletion of KIF1A and KIF5A alters the neurogenic potential of human iPSC-derived cortical progenitors, consistent with the results we obtained in *Drosophila* MNs. Single shRNAs for just KIF1A or just KIF5A did not result in a decrease of the size of neuronal clumps, nor in a reduction in the area covered by neuronal processes, which further supports some redundancy of these two motors in neurons (Supplementary Figure S3A-B).

As iPSC-derived cortical neurons depleted for both KIF1A and KIF5A seem unable to develop and differentiate as efficiently as control cells, and as axonal transport is linked to neuronal health, we hypothesised that over-expressing these microtubule motors in degenerating neurons may rescue some of their pathological defects. To test this, we analysed whether increased kinesin levels might improve survival in neuronal cells originated from patients affected from familial AD, and whether increased kinesin levels could reduce the toxicity of tau (a key component of the AD neurofibrillary tangles) and Aβ. To do so, we generated neurons starting from iPSCs derived from Down’s syndrome individuals (DSiPS). Down’s syndrome is a genetic condition resulting from having three copies of chromosome 21. Since APP is localised on the chromosome 21, this results in having three copies of the APP gene^34^ commonly linked with AD-like dementia^35^ and DSiPS neurons reproduce features of AD such as increased Aβ levels and the presence of toxic fragments of extracellular tau^36–38^. Results showed that overexpression of both KIF1A or KIF5A in DSiPS neurons (obtained by lentivirus infection) did not decrease the levels of tau (Supplementary Figure S4A). In addition, β42/β40 and β38/β40 ratios were measured in the media from the KIF1A or KIF5A overexpressing DSiPS cells, and no difference was observed compared to ratios measured in DSiPS cells (Supplementary Figure S4B).

### A**β**_42Arc_ expression affects axonal transport of mitochondria in *Drosophila*

To investigate the hypothesis that over-expressing these kinesins in degenerating neurons may rescue some of their pathological defects in *Drosophila*, we first analysed pathological defects in the Aβ_42Arc_ fly model that may be linked to kinesin function, such as the transport of mitochondria. Previous *Drosophila* work demonstrated that over-expression of hAPP and hBACE, or Aβ_42_ (wildtype or Arc) alters mitochondria localisation in MNs^14,39,40^. To explore whether the APP mutation Aβ_42Arc_ affects mitochondria axonal transport, we generated transgenic flies expressing both Mito-GFP and the human Aβ_42Arc_ peptide under the control of the *ccap-Gal4* driver, as a mean to mis-express in a small number of neurons, including a few efferent axons that could be readily imaged in peripheral nerves. This genetic combination allowed us to do precise imaging of mitochondrial transport and distribution in a single axon per peripheral segmental nerve *in vivo* in L3 larvae, an ideal developmental stage for transport studies^19,41^ (Figure 3, see materials and methods for details). To test whether the Aβ_42Arc_ expression affects the total number of mitochondria present in the segmental nerves, we quantified the total number of mitochondria using the kymograph generated from individual time points taken from our movies. We selected segmental nerves where we could easily monitor Mito-GFP, starting from the proximity of the ventral nerve cord (VNC) to the distal regions (Figure 3A). We did not observe a significant difference in the average total number of mitochondria per ROI in Aβ_42Arc_ expressing axons (8.1±0.8, n=21) compared with control axons (expressing only the Mito-GFP, 9.3±0.6, n=24) (Figure 3B). This result confirms that Aβ_42Arc_ expression did not cause a reduction in mitochondrial number in the segmental nerves.

**Figure 3.**
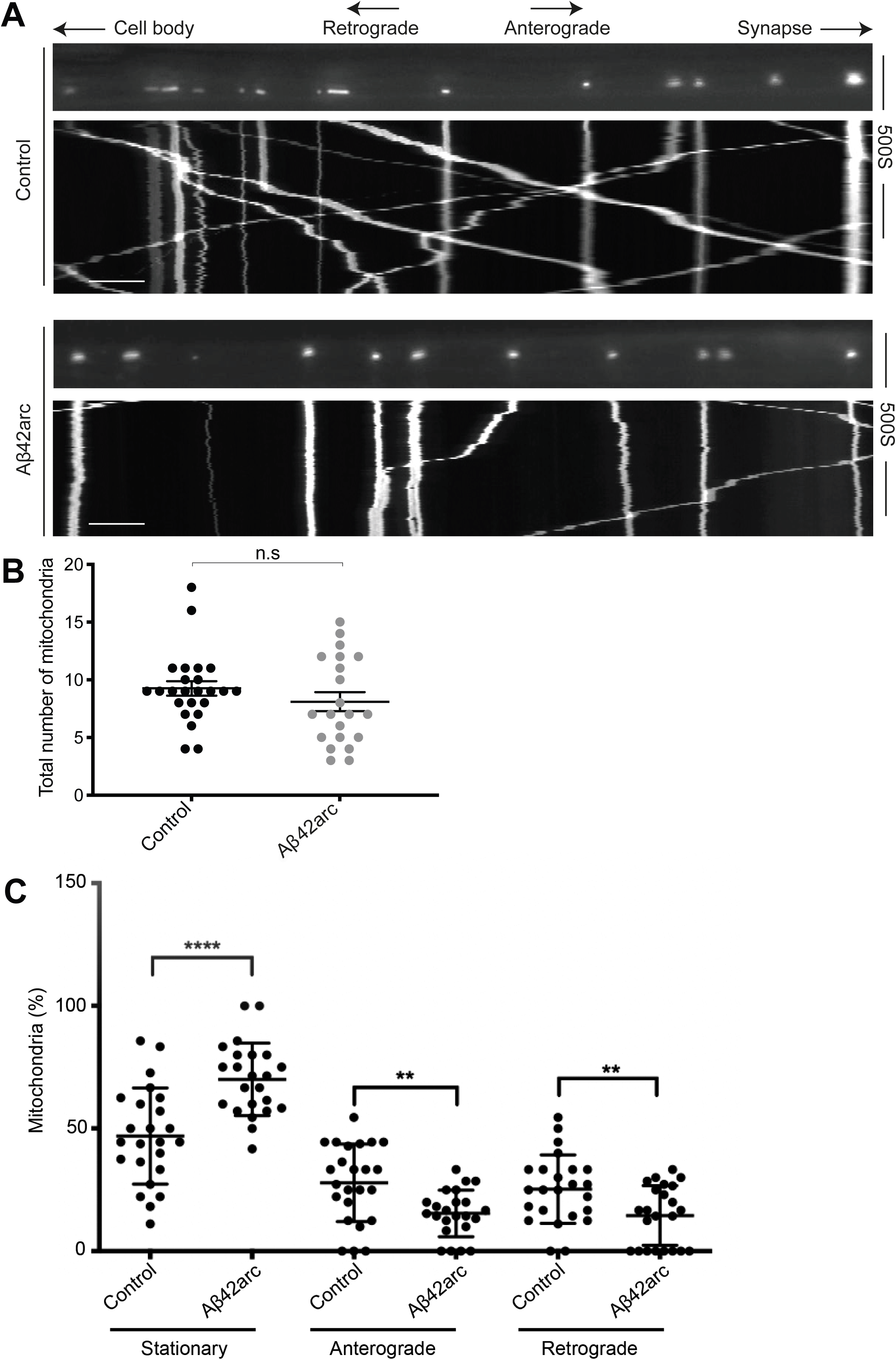
Aβ_42Arc_ expression affects axonal transport of mitochondria. (A) Representative kymograph showing mitochondrial transport in control (top) and Aβ_42Arc_-expressing (bottom) motor axons in L3 larvae during 500s time-series. GFP-labelled mitochondria were observed in motor neurons using *CCAP-Gal4, UAS-Mito-GFP* larvae. Three different categories of mitochondria were scored, i.e. stationary, anterograde, and retrograde (the direction of movement in indicated at the top). Kymographs show a significant increase in the stationary status of mitochondria, as well as a decrease in the anterograde and retrograde fractions of mitochondria, in larvae where motor neurons express human Aβ_42Arc_ compared with the control. scale bar, 10 μm. (B) Histogram showing quantification of the total number of mitochondria. (C) Histogram showing quantification of the percentage of stationery, retrograde or anterograde mitochondria populations in motor neurons from controls (n=24) and from larvae with Aβ_42Arc_-expressing motor neurons (n=23). No overall difference was observed in the total number of mitochondria between the two conditions. However, the percentage of stationary mitochondria was significantly higher, and the fraction of mitochondria in anterograde and retrograde movement was significantly lower, in Aβ_42Arc_-expressing motor neurons compared with the control. * indicates *p*<0.05. Lines show mean ± SEM.

We then quantified mitochondria dynamics in control (n=24) and in the Aβ_42Arc_ expressing neurons (n=23, and Aβ_42Arc_ neurons from here on). We were able to distinguish three discrete categories of mitochondria based on their movement - anterograde, retrograde, and stationary (Figure 3A) - and we quantified the proportions of mitochondria moving along the axons (Figure 3C). These experiments allowed us to detect that neurons from the Aβ_42Arc_ mis-expression mutants showed a significant increase in the stationary fraction of mitochondria (control: 47±4%; Aβ_42Arc_: 70±3%, *p*<0.00001), as well as a decrease in both the anterograde (control: 28±3%; Aβ_42Arc_: 15.4±2%, *p*<0.001) and retrograde (control: 25±3%; Aβ_42Arc_: 15±3%, *p*<0.001) fractions, as compared to controls (Figure 3C). This result indicates that expression of a pathologic form of human APP inhibits mitochondrial transport in *Drosophila* MNs.

We then sought to verify whether the increased number of stationary mitochondria observed in axons of Aβ_42Arc_ neurons was related to a drop in the speed of the moving organelles in these MNs. To quantify this, we tracked moving mitochondria and calculated their average velocity. We did not detect any significant difference in the mitochondrial velocity in the Aβ_42Arc_ expressing axons (anterograde, 0.9±0.1 μm/s, n=7 movies; retrograde 1.2±0.2 μm/s, n=7 movies) compared to controls (anterograde, 0.9±0.1 μm/s, n=5 movies; retrograde, 0.8±0.1 μm/s, n=5 movies). This result indicates that Aβ_42Arc_ expression did not cause a significant change in the average velocity of anterograde and retrograde mitochondria in the segmental nerves, although there is a tendency towards higher velocity in the retrograde fraction of mitochondria. This finding suggests that although the fraction of moving mitochondria in Aβ_42Arc_ axons is reduced, those mitochondria that move do it at the normal rate.

### A**β**_42Arc_ expression in MNs results in an increased number of Type Ib boutons

Several lines of evidence indicate that synapse dysfunction is part of the cellular basis of cognitive defects in AD^1,14,42,43^. We explored whether the link between Aβ_42Arc_ expression and mitochondrial transport had any effect on synapse development in L3 larvae, an ideal system to also study synaptic features^39,44^. We analysed synapse formation using the NMJs of segment A3 muscle 6/7 (NMJ6/7 from hereon), which is a common and well-established system for the study of synapse formation and is exclusively innervated by Type I boutons^45^ (Figure 4). We expressed Aβ_42Arc_ specifically in the MNs under the control of a Gal4 driver for glutamatergic neurons, including all MNs (OK371-Gal4) and we studied the morphology of the NMJ by confocal microscopy. We observed an increase in the average number of total boutons in Aβ_42Arc_ larvae (148±6, n=43) compared to the control OK371-Gal4 larvae (101±3, n=21; *p*<0.0001) Figure 4A and B). Type I boutons are subdivided into Type I big and small (Ib and Is) boutons, differing in size, morphology, physiology, and the amount of subsynaptic reticulum that surrounds them^39,46,47^. Aβ_42Arc_ larvae showed a significant increase in Type Ib boutons (37±2.6, n=20) compared to controls (29±2, n=20; *p*<0.05; Figure 5, left). However, we did not observe any significant difference in Type Is boutons (Aβ_42Arc_: 72±4, n=20 and control: 67±3, n=20; Figure 5, right). This finding is consistent with previous work reporting a significant increase in the number of Type Ib boutons in APP and APPL (*Drosophila* APP homologue) over-expressing neurons^39^. We then studied the presence of Bruchpilot (Brp), a protein specifically localised to the presynaptic release sites where synaptic vesicles fuse to the presynaptic membrane. We did not observe any significant difference in the total number of Brp puncta per NMJ from Aβ_42Arc_ larvae (604±22, n= 14) compared to the control (673±3, n=18) (Supplementary Figure S5A-B), consistent with the APP and BACE over-expression *Drosophila* models. Our results show that there is a significant increase in the number of boutons in MNs expressing the human Aβ_42Arc_.

**Figure 4.**
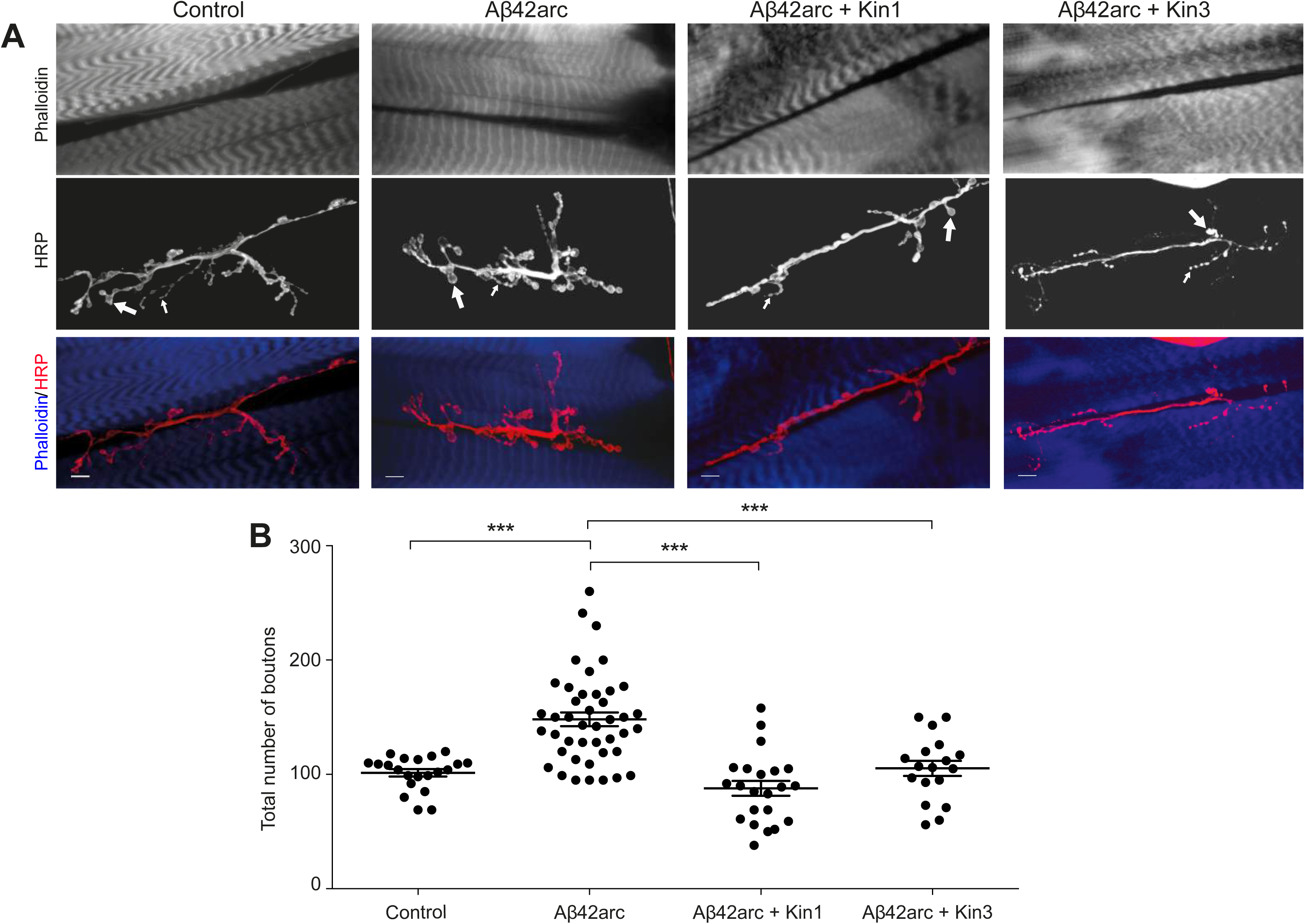
Synaptic morphology and synaptic bouton quantification in control neurons and in neurons with Aβ_42Arc_ with or without kinesin overexpression. (A) Confocal images of the NMJ at muscles 6/7 (segment A3, L3 larval stage) in controls, Aβ_42Arc_-expressing larvae, and in larvae co-expressing both Aβ_42Arc_ and either Kin1 or Kin3, stained with phalloidin (muscle staining, top row and in blue in the bottom row), and with the presynaptic neuronal membrane marker anti-HRP (middle row, and in red in the bottom row). The large and small arrows indicate Type 1b and Type 1s boutons, respectively. Scale bar, 10 μm. (B) Histograms showing the total number of boutons in the same genetic background as described in (A). Lines represent mean ± SEM, * indicates *p*<0.05.

**Figure 5.**
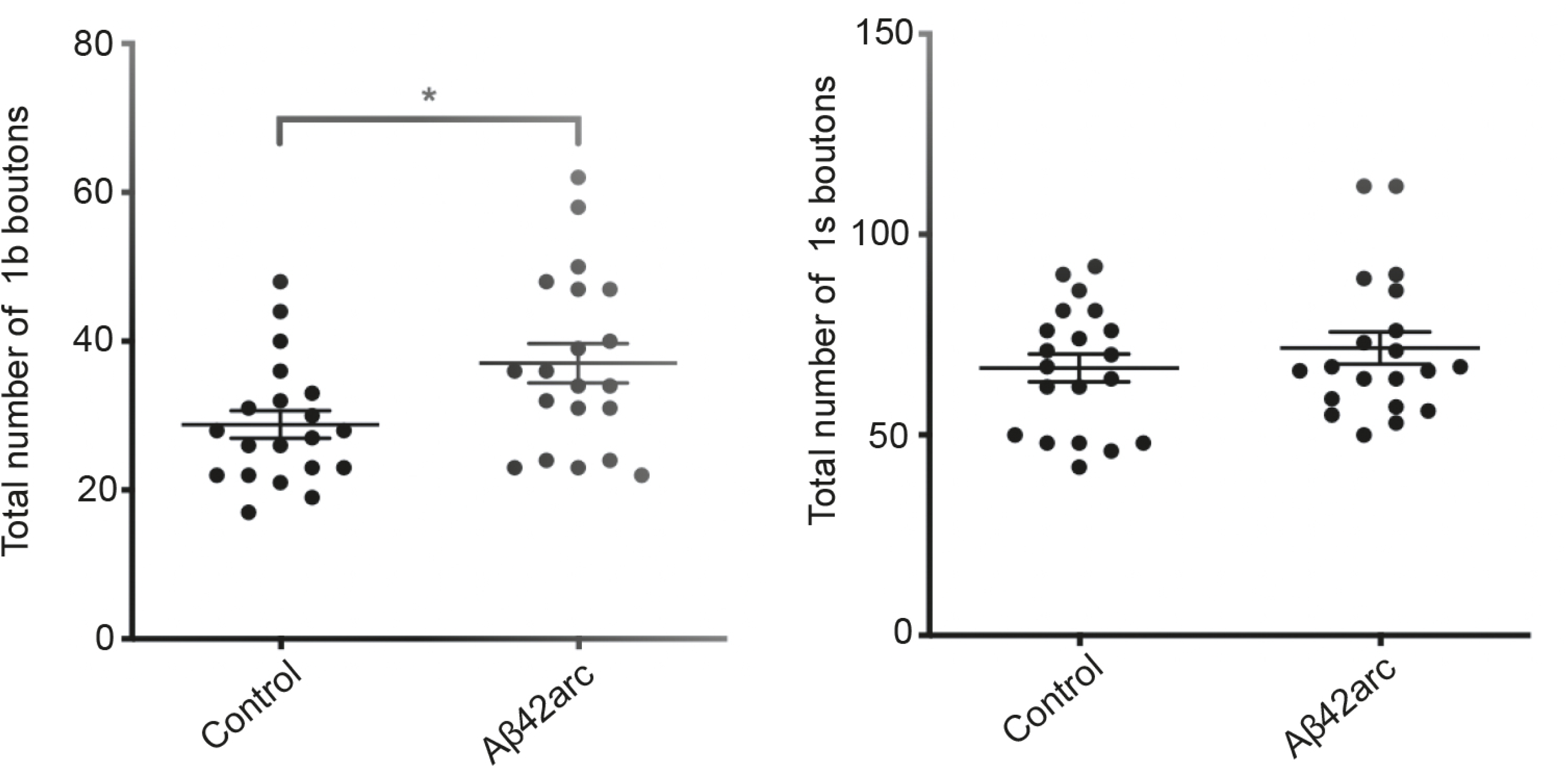
Quantification of 1b and 1s boutons in control and Aβ_42Arc_ expressing neurons. Histograms showing (left) the total number of 1b boutons and (right) the total number of 1s boutons, as quantified at NMJ 6/7 (segment A3, L3 larval stage) in controls (n=21) and in Aβ_42Arc_-expressing larvae (n=43). Lines represent mean ± SEM, * indicates *p*<0.05.

### kin1 or kin3 over-expression rescues the higher bouton number observed in A**β**_42Arc_ expressing MNs

To investigate whether the increased number of boutons observed in Aβ_42Arc_ NMJ6/7 is in any way related to kinesin activity and axonal transport, we analysed whether kin1 or kin3 over-expression might rescue this bouton specific Aβ_42Arc_ phenotype. To analyse this, kin1 or kin3 were over-expressed in conjunction with Aβ_42Arc_ in the larval MNs by using the OK371-Gal4 driver (Figure 4; see materials and methods for details). We studied the synaptic bouton numbers at NMJ6/7, and found that both kin1 (88±6, n=22) and kin3 (105±6.6, n=18) over-expression significantly suppressed the Aβ_42Arc_ induced bouton increase (148±6, n=43) (Figure 4B).

Interestingly, we noticed that the average number of boutons present in the kin1 or kin3 over-expressing larvae was similar to the number of boutons in the control (101±3, n=21, Figure 4B). We previously found no clear rescue of mitochondria transport by over-expressing kin1 or kin3. Thus, the restoration of NMJ6/7 boutons number upon higher levels of kin1 and kin3 seems not to be linked to mitochondria transport. However, it is important to keep in mind that the experiments for imaging mitochondria within a single axon *versus* the bouton rescue experiments described above required two different Gal4 drivers, which may result in different levels of expression of the motor proteins.

### Over-expression of kin1 or kin3 rescues locomotion defects in A**β**_42Arc_ *Drosophila*

To further investigate the relationship between Aβ toxicity and kinesins, we analysed how increased levels of kin1 and kin3 would impact the neurodegeneration-linked behaviour observed in our Aβ toxicity fly model. To test this, we drove the expression of Aβ_42Arc_ under the control of the pan-neuronal driver Elav-Gal4, which has been shown to recapitulate some of the AD pathologies. We chose the well-assessed method of negative geotaxis^48^ - the ability of the flies to climb against gravity - to verify age-related neurodegeneration. Analysis of the climbing capacity of the Aβ_42Arc_ flies conducted for a period of 20 days (5-25 days old flies) revealed progressive locomotion defects, with a clear drop in climbing capacity already at day 15 (Figure 6).

**Figure 6.**
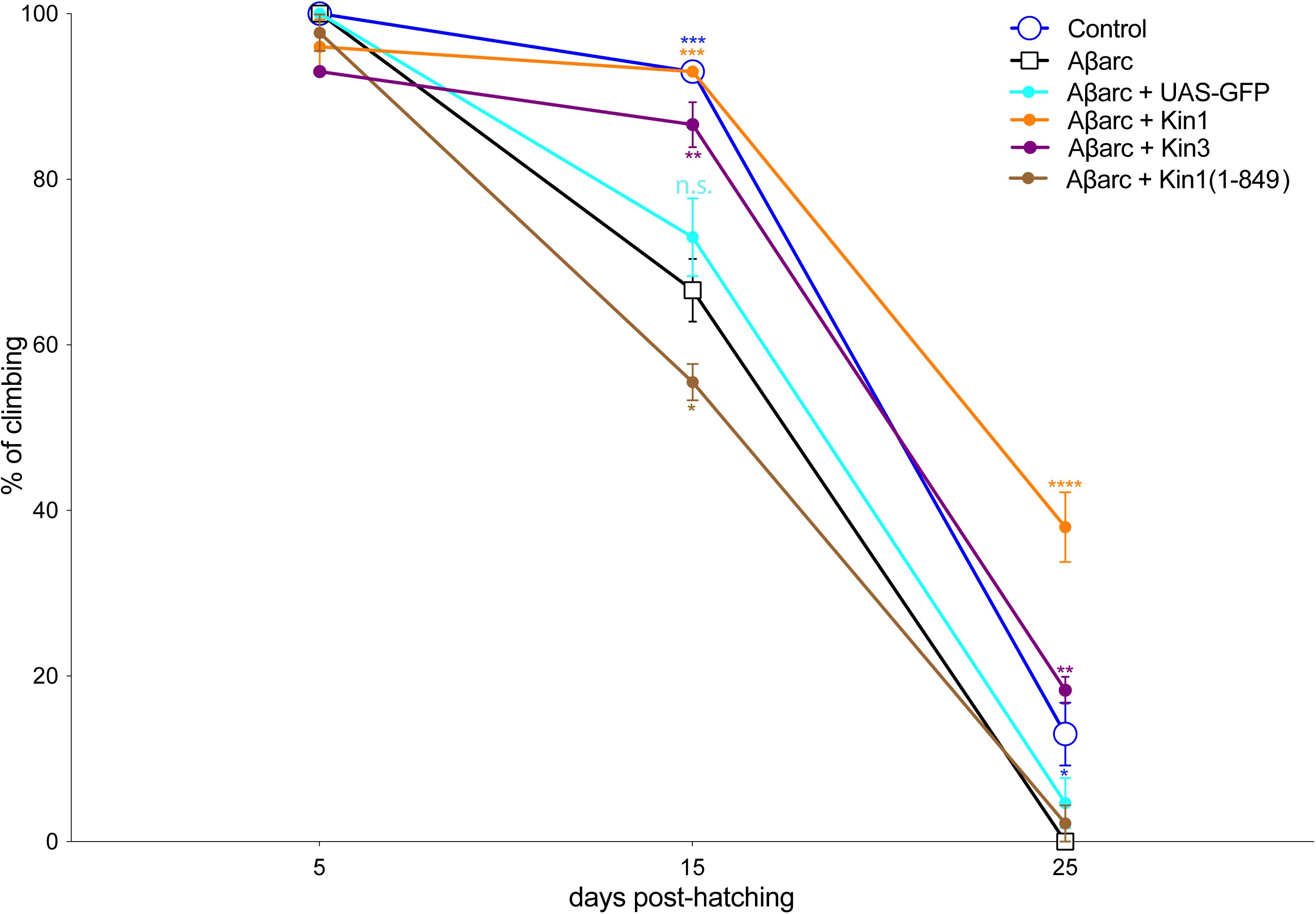
Over-expression of Kinesin-1 or Kinesin-3 rescues climbing performance of Aβ_42Arc_ *Drosophila*. Graphical profile showing the climbing performance of the flies carrying the following genotypes over a period of 0-25 days post-eclosion: Control (*elavGal4*), Aβ_Arc_ (*elavGal4; UAS-A*β*_42Arc_*), Aβ_Arc_ + GFP (*elavGal4; UAS-A*β*_42Arc_; UAS-GFP*), Aβ_Arc_ + Kin1 (*elavGal4; UAS-A*β*_42Arc_; UAS-Kin1*), Aβ_Arc_ + Kin3 (*elavGal4; UAS-A*β*_42Arc_; UAS-Kin3*), Aβ_Arc_ + Kin1(1-849) (*elavGal4; UAS-A*β*_42Arc_; UAS- Khc-849*). The X-axis shows the age of the flies (in number of days post-eclosion) and the Y-axis shows the average percentage of flies that climbed over the threshold line (see materials and methods). Climbing performance was measured from day 5 in all flies. p-values are presented in comparison with the values obtained for the overexpression of Aβ_42Arc_ and are as follows: At 15-days of age: Aβ_42Arc_ vs control and Aβ_42Arc_ vs Aβ_42Arc_ + Kin-1 ***; Aβ_42Arc_ vs Aβ_42Arc_ + Kin-3 **, Aβ_42Arc_ vs Aβ_42Arc_ + Kin1(1-849) *, Aβ_42Arc_ vs Aβ_42Arc_ + GFP n.s. At 25 days of age: Aβ_42Arc_ vs control *, Aβ_42Arc_ vs Aβ_42Arc_ + Kin-1 ****, Aβ_42Arc_ vs Aβ_42Arc_ + Kin-3 **, Aβ_42Arc_ vs Aβ_42Arc_ + GFP and Aβ_42Arc_ vs Aβ_42Arc_ + Kin1(1-849) n.s. Four experimental repeats per genotype with 15 flies per trial. *, **, *** and **** indicate p<0.05, <0.01, <0.001 and <0.0001 respectively. The error bar represents SEM.

We then generated flies over-expressing kin1 or kin3 in the same neurons that expressed Aβ_42Arc_ and analysed the climbing ability of these flies. At day 15 we already observed a significant rescue in climbing ability when either kin1 or kin3 were over-expressed in the Aβ_42Arc_ neurons, compared to controls (Figure 6). Flies that over-expressed kin1 and Aβ_42Arc_ performed as control until 15 days of age, and then performed with a significantly higher performance index than Aβ_42Arc_ flies at 25 days (Figure 6). Similar results were observed when kin3 was over-expressed in neurons of the Aβ_42Arc_ flies, with a significant rescue of climbing defects from day 15 (Figure 6). Contrary to the over-expression of the two motors, the expression of just GFP (UAS-GFP) did not rescue the climbing performance defects when co-expressed with Aβ_42Arc_ (Figure 6). This is in accordance with previous reports on another UAS construct (UAS-Glut) that does not alter Aβ_42Arc_ protein or mRNA levels compared to controls^49^. Contrary to the expression of the full length kin1, the expression of Khc1-849 (kin1 motor subunit Khc lacking the last 850-975 amino acids) did not rescue the climbing performance defects when co-expressed with Aβ_42Arc_ (Figure 6). Moreover, the expression of Khc1-849 seemed to reduce the climbing performance index of the Aβ_42Arc_ flies at 15 days (Aβ_42Arc_ vs Aβ_42Arc_ + Kin1-849 *). This suggests that kin1 requires its tail, cargo-binding region to rescue negative geotaxis in this Aβ_42Arc_ AD fly model. These results suggest that an increase in kin1 or kin3 levels can reduce the neurodegeneration that the human Aβ_42Arc_ induced in *Drosophila*.

## Discussion

Due to the length of axons, microtubule based axonal transport is crucial in maintaining the required supply of cargoes from soma to terminals (anterograde), and from terminals to soma (retrograde). This transport is also likely to be required for clearance of aggregates to maintain neuronal health. Here we show that major transport microtubule motors KIF1A and KIF5A are expressed in human iPSC-derived neurons and are found in processes with punctate pattern suggestive of cargoes. We also show that these kin1 and kin3 motors are essential for survival of MNs both in *Drosophila* and in human iPSC-derived neurons. Such essential function of these motors correlates well with the fact that mutations in KIF5A can cause hereditary spastic paraplegia^50,51^, neonatal intractable myoclonus^52^, axonal Charcot Marie Tooth disease^53^ or amyotrophic lateral sclerosis^54^, while KIF1A variants are linked to a wide range of neurodegenerative disorders^55,56^. In animal models, KIF1A knockout (KO) mice die perinatally and have defects in the transport of synaptic vesicle precursors^57^, while KIF1A-haploinsufficient animals suffer from sensory neuropathy and reduced TrkA neurons^58^. In zebrafish, peripheral axons have defective mitochondrial transport and degenerate in KIF5A mutants^59^, while a KIF5A conditional KO mouse display sensory neuron degeneration and seizures, and die soon after birth^60,61^. In *Drosophila*, *kin1* mutant larvae accumulate axonal cargos in “traffic jams” in the peripheral nerves, which could contribute to neuronal death, as we observed here, and mutations in *Drosophila* kin3 result in neuronal atrophy^62^.

From a more technical point of view, our MARCM approach offered a unique opportunity to study kinesin function in developing neurons using null alleles of kin1 and kin3, in an otherwise heterozygous animal. Since kinesin homozygous mutants are lethal in mice, zebrafish, and *Drosophila*, most previous studies have been conducted in either heterozygous individuals or hypomorphs^59,63,64^. Our approach allows to create null mutations of kin1 and kin3 in the dividing neuronal populations in a manner that the cell body, axon, dendrite or VNC lacking these molecular motors can be marked and easily identified, allowing us to follow viability of these mutant neurons *in vivo*.

To investigate further whether the neuronal death we observed in *kin1* or *kin3* mutant larvae is due to the absence of kinesins, we studied whether we could rescue this mutant phenotype by overexpression of Kin1 or Kin3. Kin1 overexpression in *kin1* mutants and Kin3 overexpression in *kin3* mutants rescued the neuronal death phenotype in larvae. Interestingly, overexpression of Kin1 in *kin3* mutants, or Kin3 in *kin1* mutants also displayed a rescue of neuronal death at the L3 stage. A possible explanation for the rescue of neuronal death observed in neurons mutant for one kinesin by overexpression of the other motor is that each kinesin may be able to take over the function of the other motor in its absence, supporting the recent findings that the concerted action of kin1 and kin3 motors promotes efficient vesicle transport^31^. This is also in agreement with the observation that the levels of Khc (motor subunit of KIF5) increased in KIF1A (kin3) homozygous mutant mice, further suggesting that kin1 motors might partially compensate for the function of KIF1A in kif1a mutants^57^. This compensation has also been suggested in zebrafish peripheral axons, where KIF5A has a role in maintenance that is only required when KIF1B is lost. Conversely, an axonal maintenance role of KIF1B is only necessary when KIF5A is reduced^59^.

Our studies using human iPSC-derived cortical neurons show that the simultaneous depletion of KIF1A and KIF5A resulted in a decrease of the size of neuronal clumps, as well as fewer cells and neurites in areas between neuronal clumps. This suggests that the neurogenic potential of the progenitors or neurite outgrowth are reduced without KIF1A and KIF5A, which indicates a conserved requirement for kin1 and kin3 in neuronal development across species, including humans. In addition, the hypothesis of a concerted and partially redundant action of kinesins in neurons is further supported by our finding in iPSC-derived neurons, where the infection of the iPSCs with lentiviral vectors encoding the shRNA for KIF1A (KIF1A^sh^) or for KIF5A (KIF5A^sh^) did not result in a decrease of the size of neuronal clumps, or in a reduction in the area covered by neuronal processes. This is in contrast with the cells expressing both KIF1A^sh^:KIF5A^sh^, which showed a dramatic decrease of the size of neuronal clumps, nor in a reduction of the area covered by neuronal processes. Although Kinesins appear to show aspects of functional redundancy for neuronal transport, and one KIF may take over the function of another KIF in its absence, previous work shows that Kinesins are not completely redundant, with distinct mutant phenotypes being described in mice and humans^65,66^. Related to this, work in zebrafish indicates that the role of KIF1B and KIF5A in peripheral sensory axon maintenance involves mitochondrial-dependent and mitochondrial-independent mechanisms^59^.

We find not only that kin1 and kin3 are essential for the survival of *Drosophila* and human neurons, but our behavioural and cell biological experiments show that increased kin1 and kin3 levels compensate for some of the inhibitory effects of expressing a pathologic form of human APP in *Drosophila*. Although the effectiveness of kinesin-mediated transport in AD still needs to be determined, there are various findings stressing that maintenance of axonal transport by upregulation of kinesins may help maintain the normal metabolism of diseased neurons, and thereby neuronal viability. Related to this, we have found that the loss of kin1 or kin3 induces neuronal death in larvae, that the expression of Aβ_42Arc_ by the pan-neuronal driver Elav-Gal4 results in reduced negative geotaxis (with age-related neurodegeneration), and that the over-expression of kinesins in the Aβ_42Arc_ neurons was able to rescue these phenotypes. Together, these results suggest that higher kinesin levels rescue Aβ_42Arc_-induced neurodegeneration. In addition, reduction of the dosage of the microtubule minus-end directed motor Dynein suppressed neuronal death caused by expressing human APP in flies, while reduction of kin1 enhanced axonal cargo accumulations in these humanised flies^67^. Also, transcriptome analysis revealed kinesin light chain (Klc)-1 splicing as an Aβ accumulation modifier in mice^68^, and the levels of various motors (KIF5A, KIF1B AND KIF21B) are increased in AD patient samples^69,70^. Together, this suggests a model in which kinesins are upregulated in AD neurons as a plastic response to the reduced activity of these motors in the diseased neurons, and thus to the reduction of cargo transport. In addition, upregulation of kinesins may ameliorate the damaging effects of intracellular protein aggregates through increased intraneuronal transport^70^. An idea that we can only speculate on is the possibility that a general increase in axonal transport, which would be achieved by increasing the activity of various kinesins, could change the biophysical properties of the axoplasm (e.g., reducing crowding and viscosity) in a way that may increase neuronal health. This is supported by our findings that overexpression of Kin1 in *kin3* mutant larvae, or Kin3 in *kin1* mutant larvae rescued neuronal death, and that increasing either Kin1 or Kin3 levels compensate for some of the inhibitory effects of expressing Aβ_42Arc_ in *Drosophila*. This agrees with reports showing that increased KIF11 expression can improve cognitive performance of an AD mouse model^71^.

The precise relationship between neurodegeneration and the function of kinesins as cargo transporters remains elusive. It is likely that the transport of specific cargoes by Kin1 and Kin3 - such as synaptic vesicles and mitochondria - plays an important role in both the essential function of these motors in neuronal survival, and in the rescue of Aβ_42Arc_ neuronal toxicity through their higher activity. This correlates both with the fact that neurons affected in AD follow a dying-back pattern of degeneration, where axonal disruption and synaptic loss precede neuronal cell death^72–74^, and with our observation that higher levels of Kin1 and Kin3 rescue the bouton number phenotype observed in Aβ_42Arc_ neurons. Expression of Aβ_42Arc_ in MNs resulted in morphologically distinct NMJs, characterised by an increase in boutons, similar to what it was reported in larvae expressing human APP or human BACE^39^, or over-expressing *Drosophila* APPL^67,75,76^. The aberrant morphology of boutons suggests that although the number of boutons is higher, they are likely dysfunctional, correlating with Aβ_42Arc_ enhancing synaptic transmission fatigue^14^. In our assay, increasing axonal transport by over-expressing kin1 and kin3 in the neurons that also express human Aβ_42Arc_ allows for these axons to develop proper bouton number and normalised morphology. This result, together with the rescue of climbing capacities of adults when kin1 and kin3 are over-expressed in Aβ_42Arc_ neurons, suggests that higher kinesin levels also rescue the functionality of the NMJ. As previously suggested, NMJ defects may lead to reduced connectivity and innervation of those neurons and their targeted muscles, which ultimately causes locomotion defects. This model fits with our finding that the overexpression of kinesins in Aβ_42Arc_ neurons rescues both bouton morphology and animal climbing capacity. However, as boutons are analysed in larvae, while negative geotaxis is used as a readout of neurodegeneration is performed in aging adults, it is difficult to conclude on the relationship between these two processes and the function of kinesins as axonal transport motors in degenerating neurons. One possibility is that kinesins rescue the reduced axonal transport of mitochondria in Aβ_42Arc_ neurons, resulting in a proper number of functional mitochondria reaching the synapses. This hypothesis aligns with the idea that Aβ probably disrupts synaptic function by affecting presynaptic mitochondria^77^. In Aβ_42Arc_ larvae, we failed to rescue reproducibly the mitochondria transport defects by overexpressing the kinesins. However, it is still possible that the over-expression of kin1 and kin3 helps the axonal transport of mitochondria in adults and this may contribute to improving the age-associated locomotor activity of Aβ_42Arc_ flies. What our results show from the mechanistical point of view is that the tail region of Kin1 is required to rescue the neuronal dysfunction observed in Aβ_42Arc_ mutants. The tail region (amino acids 850-975 in *Drosophila)* is a conserved region and acts as an alternative cargo binding domain to the Klc cargo-binding domain in oocytes^78,79^, in mitochondria axonal transport^80^, and in mitochondria density in sensory neurons^59^.

This work sheds light into the association between the activity of the major microtubule transport motors kin1 and kin3 and Aβ neuronal toxicity. It could be argued that novel AD therapeutic strategies based on enhancing the activity of microtubule anterograde motors should be pursued. While it was previously shown that reducing KIF5B ameliorates the phenotypes in a tauopathy mouse model^81^, excess of motor activity, however, can lead to transport defects^79^. Hence, it appears that maintaining the proper balance of kinesin activity is crucial for neuronal health and may aid in mitigating AD pathogenesis.

## Materials and Methods

### Resources table

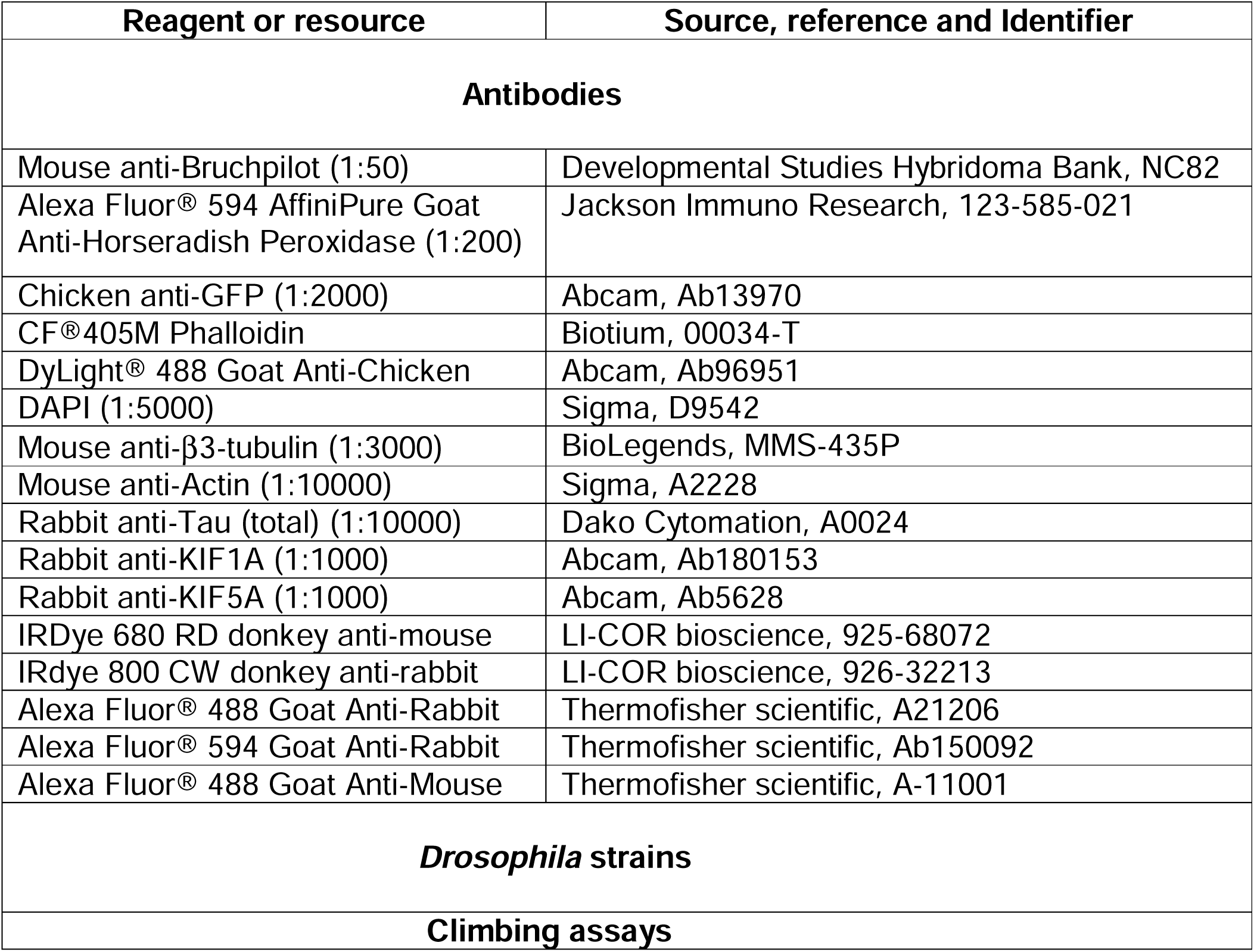
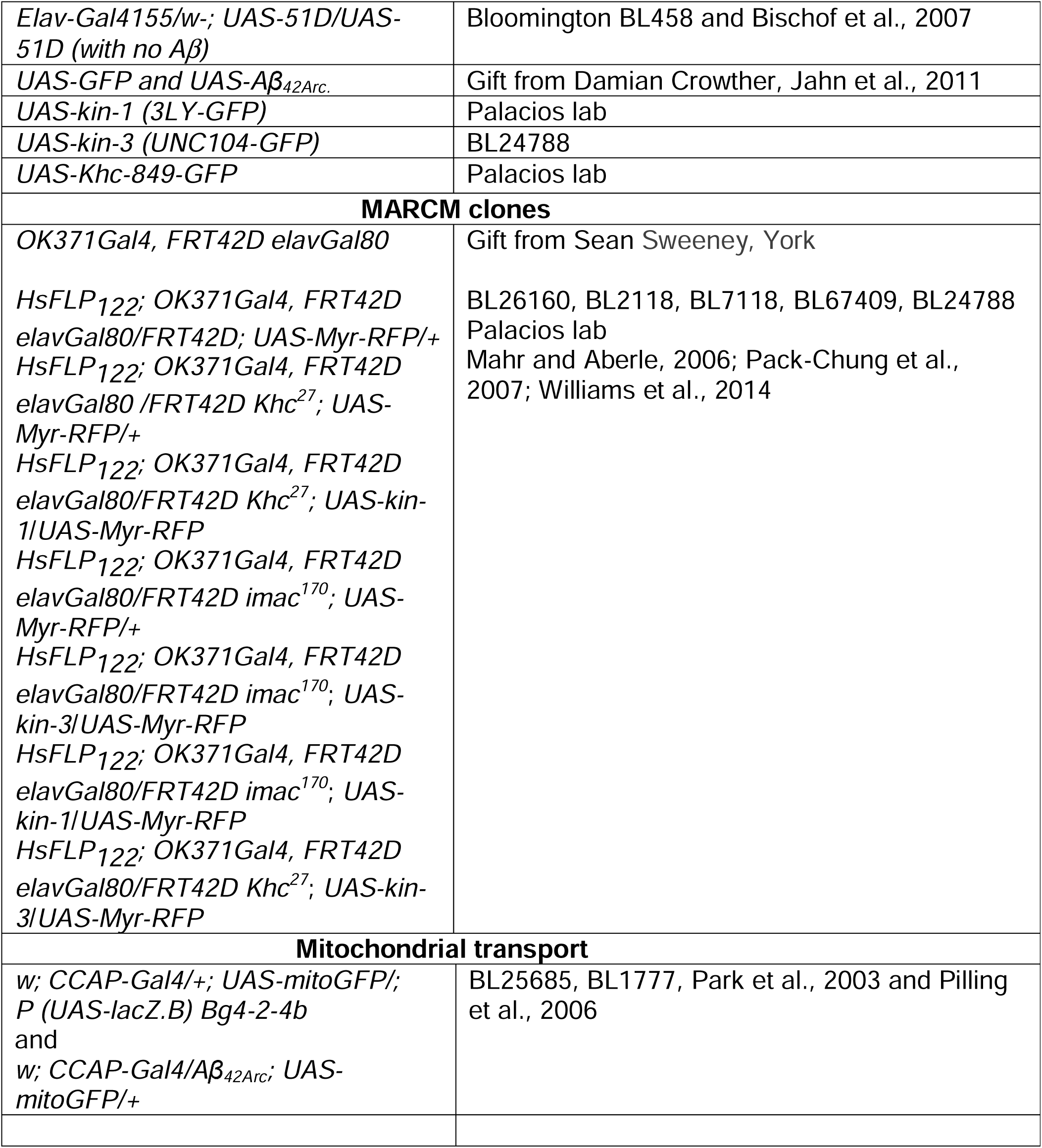

### Induction of MARCM clones, live imaging, and analysis of neuronal survival

All the flies used for the MARCM experiments were maintained at 25°C in normal fly food vials containing agar, cornmeal, molasses and yeast. Flies of the desired genotype were allowed to lay their eggs on yeast spread apple juice plates over the course of five-hour intervals. Plates were heat-shocked for 30 minutes in a 37°C water bath directly after collection. This protocol maximises clonal induction in neurons, while avoiding clonal induction in muscles. After 24 hours, the hatched L1 larvae were collected from the plate and aged for 24 more hours to reach L2. The larvae with the right genotype were selected based on markers and fluorescence^82^. Larvae with clones in the MNs (expression was driven by OK371-Gal4) were selected by the presence of myr-RFP (the myristoyl group targets proteins to membranes) in the neurons. FRT control, *kin1* or *kin3* mutants were analysed in parallel. Plates with eggs were kept in a humidified box to prevent drying. A few hours before imaging, we allowed the larvae to crawl on the apple juice plate to get rid of the yeast from the gut and the body (to prevent auto-fluorescence). For imaging, one larva per slide (L2 and L3) was mounted with 1x PBS. The cover slip was pressed gently to avoid killing the larvae and to prevent excessive movement. After imaging the L2 larvae, we put the larvae back on to the apple juice plate, so they grew until the L3 stage. The expression pattern of clones in the L2 and L3 larvae were recorded using a Zeiss Axiophot (Axioskop-40) widefield fluorescence microscope fitted with an AxioCam MR operated by the AxioVision software (Zeiss). Images were processed using ImageJ (NIH) and Photoshop (Adobe) softwares. For statistics, we used One-Way ANOVA with multiple comparison tests between the genotypes and these tests were done using Prism 7 (GraphPad).

### Generation of iPSC-derived cortical neurons

Human iPSCs were differentiated into cortical neurons as previously described^32^. Briefly, iPSCs were plated on Geltrex (Thermo Fisher Scientific)-coated plates to reach full confluence. One day after seeding (Day 0), neural induction was initiated by changing the culture medium to a 1:1 mixture of DMEM/N-2 (DMEM/F-12 + GlutaMAX (Life Technologies, cat# 31331-028); 1X N-2 (Life Technologies, cat# 17502-048); 5 μg mL^-1^ insulin; 1 mM L-glutamine; 100 μM non-essential amino acids; 100 μM β-mercaptoethanol; 50 U mL^-1^ penicillin and 50 mg mL^-1^ streptomycin) and Neurobasal/B-27 (Neurobasal (Life Technologies, cat# 12348-017); 1X B-27 (Life Technologies, cat# 17504-044); 200 mM L-glutamine; 50 U mL^-1^ penicillin and 50 mg mL^-1^ streptomycin) media (hereafter referred as N2B27) supplemented with 1 μM dorsomorphin and 10 μM SB431542 to inhibit TGFβ signalling and support neuronal differentiation and neurogenesis, N2B27 media was replaced every 24 hours. At day 12 the obtained neuroepithelial sheet was harvested and dissociated using the enzyme Dispase (Life Technologies, cat# 17105) and plated on laminin-coated plates. After one day, media was changed to N2B27 containing 20 ng/mL FGF2. N2B27+FGF2 was added freshly daily for 4 days to promote the maturation of neural rosettes. After 4 days FGF2 was withdrawn, and neural rosettes were maintained in N2B27 refreshing medium every other day. At day 30 neural rosettes were dissociated using Accutase (Innovative Cell Technologies, cat# AT104) and neural progenitor cells were plated on laminin-coated plates at 150,000 cells/mm^2^. Plated neurons were maintained for up to 120 days with a medium change every other day.

### Immunohistochemistry, confocal imaging and image analysis of iPSC-derived neurons

For confocal analysis, cells were seeded at 150,000 cells/mm^2^ in CellCarrier-96 Ultra Microplates (Perkin Elmer) and cultured until day 70. Immunofluorescent staining was performed as follow. Cells were washed 3 times in PBS and then fixed using 4% paraformaldehyde (v/v) in PBS for 15 minutes at RT. After 3 washes in PBS, cells were permeabilised in PBS+0.3% Triton X-100 (Sigma; Tx) for 15 minutes at room temperature (RT). After 3 washes in PBS, cells incubated for 1 h at RT in 5% BSA (Sigma) (w/v) in PBS+0.3% Triton X-100 (PBS-Tx+5% BSA). Primary antibodies were diluted in PBS-Tx+5% BSA and incubated overnight at 4°C. Cells were washed 3 times in PBS and incubated 1 hr in the dark at RT with secondary antibodies diluted 1:1000 in PBS-Tx+5% BSA. After 3 washes in PBS, samples were incubated for 5 minutes at RT with DAPI diluted 1:5000 in PBS and then washed 3 additional times with PBS. Cells were then left in 200 μL of 1X PBS for imaging. Confocal images were obtained using an Olympus Inverted FV3000 confocal (Olympus Scientific solutions) and processed using Fiji software^83^. For neurites extension analysis, plates were imaged using an Opera Phenix High-Content screening system (Perkin Elmer) and images analysed using the built-in Harmony software. Antibodies against KIF1A (ab180153) and KIF5A (ab5628) were obtained from Abcam and used at 1:1000 dilution. β3-tubulin (MMS-435P) was obtained from BioLegends and used at 1:3000 dilution.

### Protein extraction and western blot analysis

Total cell protein was extracted using RIPA buffer (Sigma) supplemented with protease inhibitors (Sigma) and Halt phosphatase inhibitors (Thermo Fisher Scientific). Protein quantification was performed using Precision Red Advanced Protein Assay buffer (Cytoskeleton, Inc.). For each sample, 30 μg of protein were mixed with 1X NuPAGE LDS Sample Buffer (Thermo Fisher Scientific) + 1 μM Dithiothreitol. Samples were heated at 100°C for 10 minutes and loaded on a NuPAGE 4%–12% Bis-Tris gel (Thermo Fisher Scientific). Afterward, proteins were wet transferred onto PVDF membrane (Millipore) for 1 h at 100 V. Membranes were blocked for another 60 min in 5% BSA in PBST (PBS containing 0.05% Tween 20). All primary antibodies were incubated overnight in 5% BSA in PBST at 4°C. Next day, membranes were incubated for 1 h in secondary antibody and washed gently in PBST buffer for further 30-60 min. Immunoblots were detected using LI-COR Odyssey CLx Infrared Imaging System and processed with the Image Studio Software (LI-COR).

### Infection of iPSC-derived neurons with MISSION shRNA or over-expression constructs

MISSION shRNAs against human KIF1A (SHCLNG-NM_004321) and KIF5A (SHCLNG-NM_004984) were obtained from Sigma-Aldrich. Scramble vector was the MISSION pLKO.1-puro non-Mammalian shRNA Control vector. Lentiviral over-expression constructs were obtained as follow. cDNA of human KIF1A (MHS6278-211690363) and human KIF5A (MHA6278-202800246) were amplified from Dharmacon library (Horizon discovery) and cloned inpCR-BluntII-TOPO and pCMV-SPORT6, respectively. cDNA was then subcloned in the lentiviral vector pCSC-SP-PW-GFP (pBOB-GFP) (Addgene) for neuron infection using the NEBuilder HiFi DNA Assembly Cloning Kit (New England Biolabs). iPSC-derived neurons were infected using viral particles diluted in N2B27 media (5 MOI) for 12h. After infection, media was replaced with fresh N2B27 and changed every 48 hours.

### Dissection, imaging and analysis of mitochondrial axonal transport

For mitochondrial studies we decided to use *ccap-Gal4* instead of *OK371-Gal4* because the *ccap-Gal4* driver is expressed only in a few neuronal cells, which secrete the ccap (crustacean cardioactive peptide) neuropeptide^84^. These neurons send out only one axon per segmental nerve. Therefore, we could do precise imaging of mitochondrial transport in single axons of the segmental nerves. To image L3 larvae, we washed them in fresh dissection solution (128 mM NaCl, 1 mM EGTA, 4 mM MgCl_2_, 2 mM KCl, 5 mM HEPES and 36 mM sucrose, pH 7.2.)^85^, opened them and immobilised them for imaging. Imaging of larval axons was performed as described in ^86^ and ^87^, and in summary: wandering third instar larvae were pinned at each end with dorsal side up to a reusable Sylgard (Sigma 761028) coated slide using pins (Fine Science Tools FST26002-10) cut to ∼5 mm and bent at 90°. The larvae were cut along the dorsal midline using micro-dissection scissors. Internal organs were removed with forceps without disturbing the ventral ganglion and MNs. Larvae were then covered in dissection solution. The cuticle was then pulled back with four additional pins. The anterior pin was adjusted to ensure axons are taut and as flat as possible for optimal image quality.

Movies were taken using a Nikon E800 microscope with a 60× water immersion lens (NA 1.0 Nikon Fluor WD 2.0) and an LED light source driven by Micromanager 1.4.22 Freeware^88^. A CMOS camera (01-OPTIMOS-F-M-16-C) was used to record 100 frames at a rate of 1 frame per 5 s (8.3 min total). Axons were imaged within 200 µm of the ventral ganglion in the proximal portion of the axons and no longer than 30 min after dissection. A minimum of 23 movies was taken for each genotype and roughly 2-3 axons imaged per larvae. Movies were converted into kymographs using Fiji^83^, and mitochondrial motility was quantified manually with the experimenter blinded to the condition. In each movie (filmed at 10 frames per second), regions of interest (120 x 10 μm, based on the length of axon in focus suitable for a movie in most dissections) were analysed from the proximity of the VNC to the distal region of the axon at each time point. The number of each fraction of mitochondria (stationary, anterograde and retrograde) was quantified and percentage of each fraction of mitochondria was calculated for each movie. Unpaired student ‘t’ test was used to determine the significance between different genotypes.

### Calculation of average number of mitochondria and average velocity of mitochondria

Average numbers of mitochondria in the axons of the control and the *Aβ_42Arc_* were calculated manually using the same movies which were used to generate the kymograph using ImageJ with the same ROI. Unpaired student ‘t’ test was used to determine the significance between different genotypes. The average velocity was calculated by determining the speed of each mitochondrion from the kymograph generated. For each mitochondrion, we calculated the first velocity until it made an obvious shift, by drawing a straight line on ImageJ (change of direction or pause represent different straight lines on a kymograph). Then the second velocity was calculated until the next drift and the process continued until it finished anteriorly or posteriorly. After calculating the velocity for each shift for each mitochondrion in a series of images, we calculated the average mean velocity for each mitochondrion and finally calculated the average velocity of both anterograde and retrograde separately. The velocity was calculated using the ‘Velocity Measurement Tool’ on ImageJ (http://dev.mri.cnrs.fr/projects/imagejmacros/wiki/Velocity_Measurement_Tool).

### Egg collection, larval staging and analysis of boutons and cargoes in NMJ of the 3^rd^ instar larvae

Males and females of the desired genotype flies were put in a cage with yeast plated apple juice plate and kept at 29°C. The flies were allowed to lay eggs on fresh apple juice plate every day for the next two days and these plates were discarded. On the third day the flies were transferred on to a fresh apple juice plate for the collection and selection of the late 3^rd^ instar larvae. Although the 3^rd^ instar can be distinguished based on the number of teeth in their mouthparts, precise staging is difficult because of the asynchronization of the development of the larvae. Wandering larvae for fillet making were staged by feeding them with thick yeast paste mixed with bromophenol blue sodium salt (sigma, B5525). The wandering larvae stop feeding and the blue dye gradually disappears from their intestine completely. Larvae with blue or white gut were considered to be early L3 (12-24 hours before pupariation) or late L3 (1-6 hours before pupariation), respectively^89,90^. We mainly selected larvae with partially cleared gut and fully cleared gut for bouton number analysis, as the new synaptic boutons are continually added until the late 3^rd^ instar larval stage. Larvae were placed in saline solution, pinned dorsal side up and dissected from posterior to anterior to obtain fillet preparation. The body was extended and pinned on both side; the gut was carefully removed, and fillet were washed with saline solution. Larvae were fixed (4% formaldehyde), transferred into a solution of PBS-Triton (0.3%) and immune-stained using the set of primary and secondary antibodies indicated in the table above. Samples were transferred in glycerol and imaged using Leica TCS SP5 upright confocal microscope; images were processed using ImageJ (NIH) and Photoshop (Adobe). All the images were acquired from the muscle 6/7 of abdominal segment A3 of the larval fillet and we manually counted the total number of boutons, synaptic puncta and Bruchpilot puncta. We used one-way ANOVA statistical tests for multiple comparison and unpaired student ‘t’ test for the comparison between two genotypes; these tests were done using Prism 7 (GraphPad).

### Negative geotaxis climbing assay

Stocks used for these experiments were isogenised by backcrossing them six times with w1118 flies. To assay the climbing ability of the control, Aβ_42-arctic_ and rescue flies, 15 female flies of the same age were collected for each genotype and placed in different 25ml plastic pipettes. We left the flies in the pipettes for at least five hours prior to the experiment, for them to acclimatize to the pipette and get rid of the CO_2_ effect^91^. Experiments were performed at a room temperature of 25°C. In each experiment, the flies were tapped down to the bottom of the plastic pipette and allowed to climb for one minute. The numbers of flies crossing the 15ml line (i.e. at around 3.8 cm from the bottom) and of those remaining at the bottom of the pipette were recorded. The assays were repeated three times with 15 flies for each genotype and for different ages. The graph in Figure 6 represents average of the results from the three independent trials and expressed as the average percentage of the total number of flies in the tube (= % of climbing activity). Statistical significance was assessed between genotypes over time by unpaired student’s t test (comparison made between two genotypes) using GraphPad Prism 7.

## Acknowledgements

We thank Sean T. Sweeney and Damian Crowther for reagents; Matthias Landgraf, Charalampos Rallis and Teresa Niccoli for manuscript comments and discussions. Matthew Oswald and Matthias Landgraf for stocks, immense help with the *Drosophila* work and for discussions.

## Author contributions statement

DF, FP, CCGF, and IMP designed and performed experiments, analysed data, and wrote the paper. VLH designed and performed experiments and analysed data. MEG and IP performed experiments and analysed data. AW contributed to conceptualisation and project supervision. FJL designed experiments and contributed to conceptualisation and project supervision.

---

**Supplementary figure S1.**
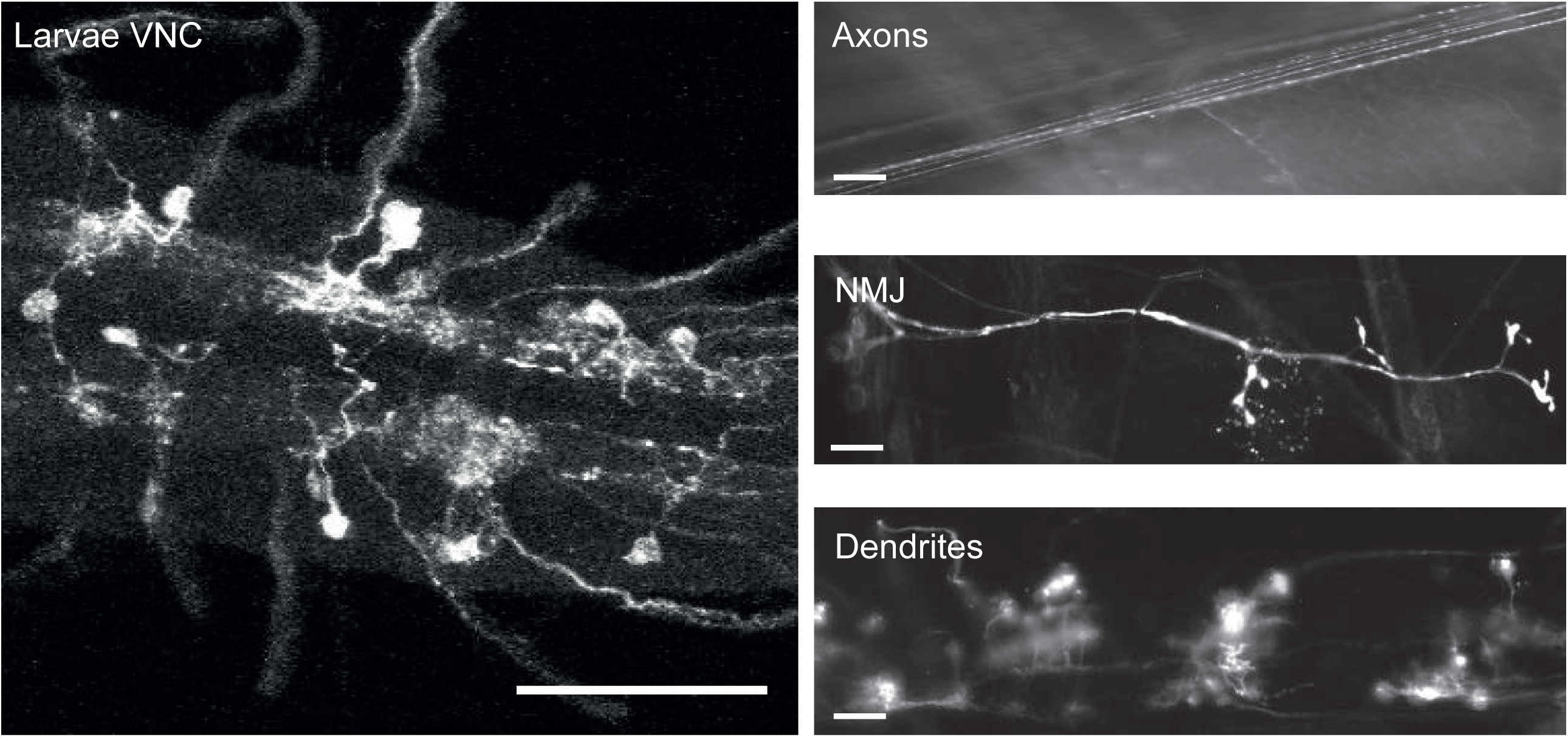
Images showing the labelling of neurons by the MARCM technique from a 3^rd^ instar larval fillet. Left: Central region of the ventral nerve cord in which some motor neurons are labelled thanks to the MARCM technique using the driver line OK371-Gal4 and the UASmyr-RFP effector. Anterior to the left. Scale 50 μm. Right: Enlarged view of motor neurons labelled with OK371Gal4 and UASmyr-RFP, with at the (top) a view of the axons, in the (middle) a view of the axons where they join the muscle fibre and form boutons at the neuromuscular junctions (NMJ 6/7), and at the (bottom) a view of the neuronal dendrites. Anterior to the top. Scale bar, 10μm.

**Supplementary figure S2.**
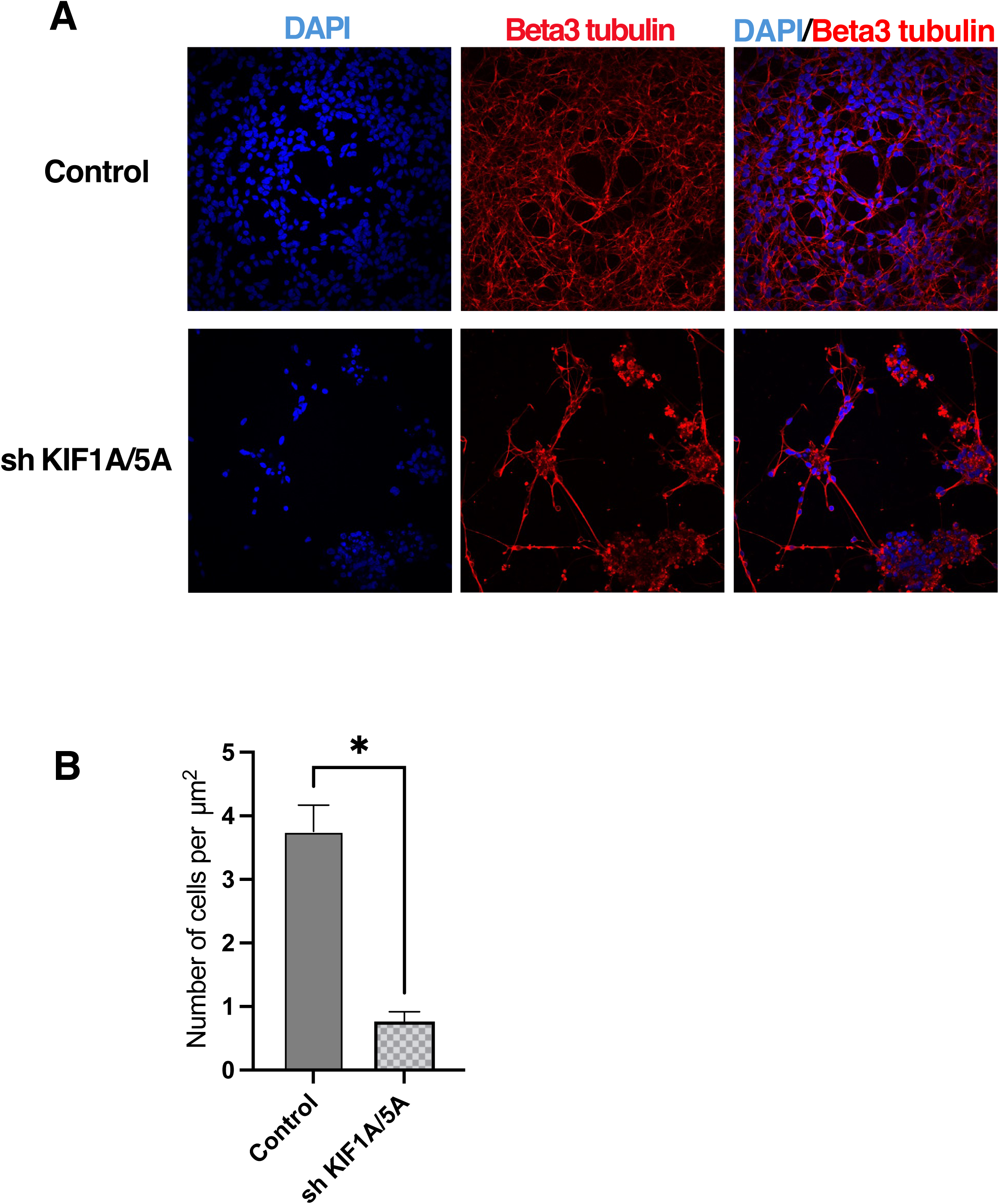
Immunostaining and cell number after silencing both KIF1A and KIF5A in iPSC-derived neurons. (A) iPSC-derived neurons were infected at day 30 with both shRNA constructs against KIF1A and KIF5A and cultured to day 65 when cells were fixed and stained for DAPI (blue) and β3-tubulin (red). (B) A comparison of the number of cells present in areas between neuronal clumps in control neuronal cultures (n=4 areas of 0.150mm^2^) and in neuronal cultures infected by KIF1A^sh^:KIF5A^sh^ (n=3 areas of 0.150mm^2^) (*p*=0.0020, t-test).

**Supplementary figure S3.**
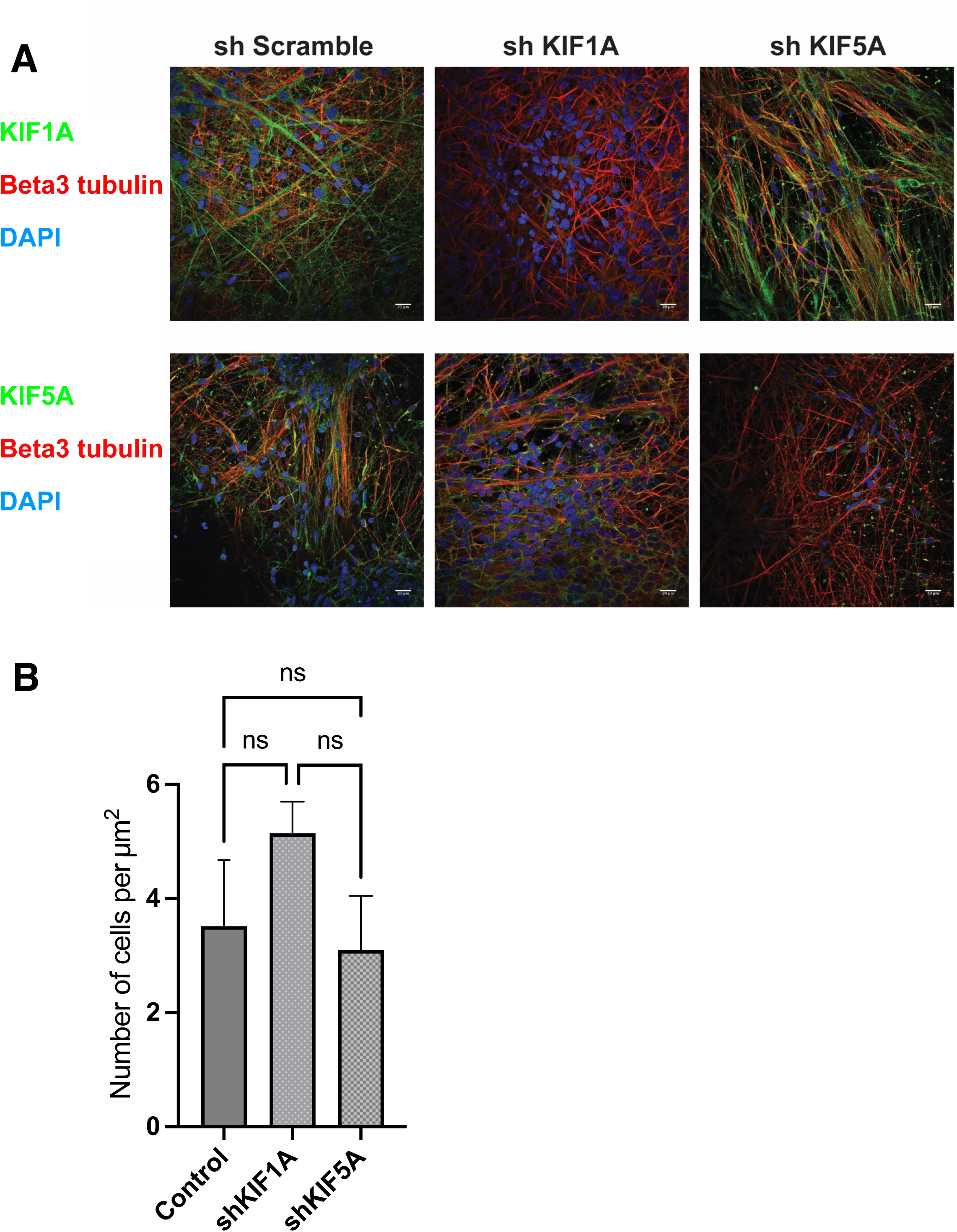
Immunostaining and cell number after silencing either KIF1A or KIF5A in iPSC-derived neurons. (A) iPSC-derived neurons were infected at day 30 with either shRNA constructs against KIF1A or KIF5A and cultured to day 65 when cells were fixed and stained for DAPI (blue) and β3-tubulin (red), as well as for either KIF1A or KIF5A (green). (B) A comparison of the number of cells present in areas between neuronal clumps in control neuronal cultures (n=3 areas of 0.150mm^2^) and in neuronal cultures infected by sh KIF1A (n=5 areas of 0.150mm2) or sh KIF5A (n=5 areas of 0.150mm^2^) (Kruskal-wallis multi-comparison test).

**Supplementary figure S4.**
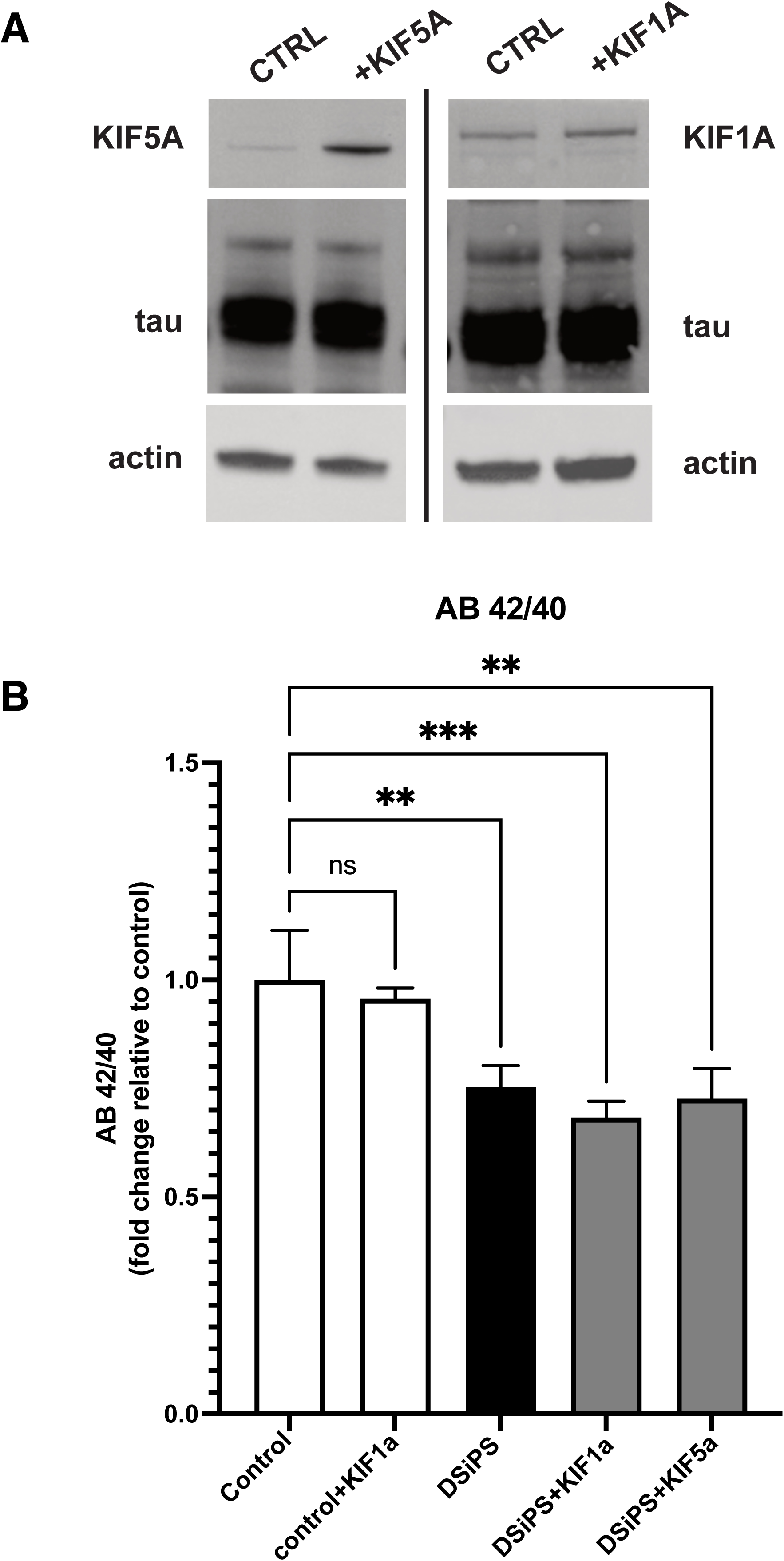
Protein extracts and Aβ42/40 ratios after overexpressing KIF1A or KIF5A in iPSC-derived DSiPS neurons. (A) Protein extracts after infection of iPSC-derived DSiPS neurons with Lentiviral over-expression constructs for KIF1A and KIF5A. Western blots show that KIF1A and KIF5A levels are increased, but that tau and actin levels were not affected in cells with higher amount of kinesins. (B) Aβ42/40 ratio was analysed in conditioned media from iPSC-derived neurons (control) and iPSC-derived DSiPS neurons. A decrease is detected in DSiPS neurons compared to control neurons. No effect on Aβ42/40 ratio was reported after over-expression of KIF1A or KIF5A.

**Supplementary figure S5.**
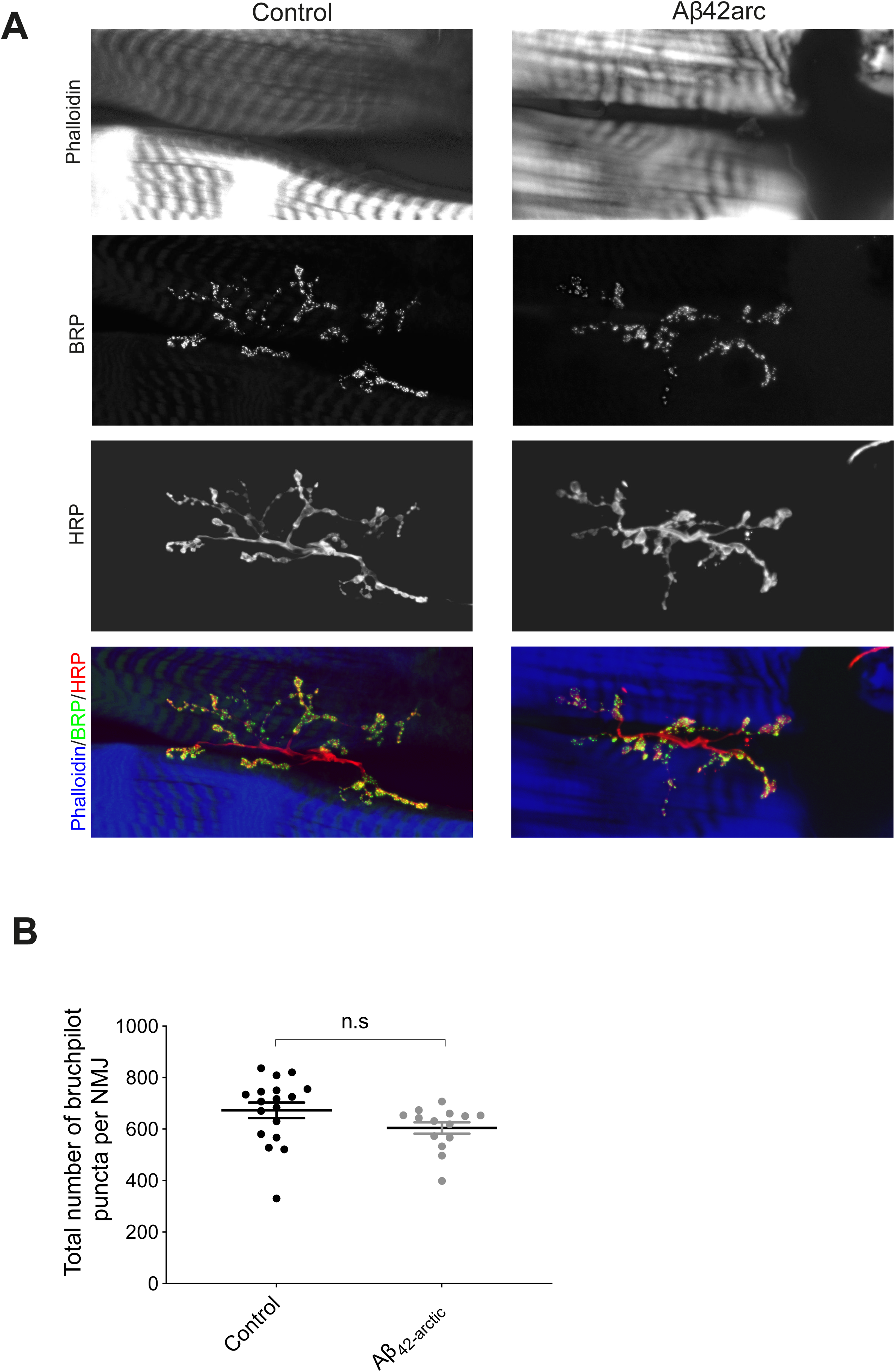
Quantification of the Bruchpilot puncta at the NMJ 6/7 of control and Aβ_42Arc_ larvae. (A) Confocal images of the NMJ 6/7 (segment A3, L3 larval stage) stained with the active zone protein marker Bruchpilot (Brp, green in the bottom row) and with the presynaptic neuronal membrane marker anti-HRP (HRP, red in the bottom row). F-actin in muscles is labelled with phalloidin (Top panel and blue in the bottom row). Scale bar, 10 μm. (B) Quantification of the Bruchpilot puncta present in the larval NMJ6/7 in control (n=18) and in Aβ_42Arc_ (n=14) larvae. No difference can be observed in the number of puncta between these two types of larvae.

